# CozEa and CozEb play overlapping and essential roles in controlling cell division in *Staphylococcus aureus*

**DOI:** 10.1101/256560

**Authors:** Gro Anita Stamsås, Ine Storaker Myrbråten, Daniel Straume, Zhian Salehian, Jan-Willem Veening, Leiv Sigve Håvarstein, Morten Kjos

**Affiliations:** Faculty of Chemistry, Biotechnology and Food Science, Norwegian University of Life Sciences, Ås, Norway.; Department of Fundamental Microbiology, Faculty of Biology and Medicine, University of Lausanne, Lausanne, Switzerland.

## Abstract

*Staphylococcus aureus* needs to control the position and timing of cell division and cell wall synthesis to maintain its spherical shape. We identified two membrane proteins, named CozEa and CozEb, which together are important for proper cell division in *S. aureus*. CozEa and CozEb are homologs of the cell elongation regulator CozE^Spn^ of *Streptococcus pneumoniae*. While *cozEa* and *cozEb* were not essential individually, the Δ*cozEa*Δ*cozEb* double mutant was lethal. To study the functions of *cozEa* and *cozEb*, we constructed a CRISPR interference (CRISPRi) system for *S. aureus*, allowing transcriptional knockdown of essential genes. CRISPRi knockdown of *cozEa* in the Δ*cozEb* strain (and vice versa) causes cell morphological defects and aberrant nucleoid staining, showing that *cozEa* and *cozEb* have overlapping functions and are important for normal cell division. We found that CozEa and CozEb interact with the cell division protein EzrA, and that EzrA-GFP mislocalizes in the absence of CozEa and CozEb. Furthermore, the CozE-EzrA interaction is conserved in *S. pneumoniae*, and cell division is mislocalized in *cozE*^Spn^-depleted *S. pneumoniae* cells. Together, our results show that CozE proteins mediate control of cell division in *S. aureus* and *S. pneumoniae*, likely via interactions with key cell division proteins such as EzrA.

## Introduction

Bacterial cell division initiates when the tubulin-like protein FtsZ polymerizes into a ring structure (Z-ring) located at the future division site. The Z-ring then serves as a scaffold for recruitment of cell division and cell wall synthesis proteins, forming the multiprotein complex known as the divisome. Several of the proteins constituting the divisome are widely conserved in most bacteria, while others are specific for bacterial subgroups or have diverged significantly (Pinho *et al.*, 2013). Positioning and timing of Z-ring assembly and cell wall synthesis are dependent on the shape of the bacterium; and there are large variations between coccal, ovococcal and rod-shaped bacteria.

*Staphylococcus aureus* often serves as the model organism for cell division studies in spherical bacteria. *S. aureus* is an opportunistic pathogen, which persistently colonizes around 20 % of the human population (Grice & Segre, 2011), causing both superficial infections on the skin and invasive, life-threatening sepsis as well as endocarditis in humans (Rasigade & Vandenesch, 2014, Foster *et al.*, 2014). Furthermore, *S. aureus* is an important pathogen among livestock (causing mastitis and other infections) and is a problematic food pathogen. Treatment of *S. aureus* infections with antibiotics is increasingly challenging due to the rise of antibiotic resistant strains, including MRSA (methicillin-resistant *S. aureus* which are resistant to β-lactam antibiotics) and VRSA (vancomycin-resistant *S. aureus*).

Cell division in spherical *S. aureus* occurs in three consecutive planes, where every new round of division is orthogonal to the previous division plane (Pinho *et al.*, 2013). Many key cell division proteins known from other model bacteria are conserved in *S. aureus*, including FtsZ, FtsA, EzrA, GpsB, DivIB, DivIC, FtsL, MurJ, DivIVA, MreC and MreD (Pinho & Errington, 2003, Pinho & Errington, 2004, Steele *et al.*, 2011, Pinho *et al.*, 2013, Bottomley *et al.*, 2014, Monteiro *et al.*, 2018). These proteins are in different ways involved in formation of the division ring and for ensuring proper cell wall synthesis and cell division. For example, EzrA is a key early cell division protein linking division ring formation and the cell wall synthesis machinery (Jorge *et al.*, 2011, Steele *et al.*, 2011). Synthesis of new cell wall in staphylococci mainly occurs at midcell. The recruitment of cell wall synthesis proteins (i.e. transpeptidases PBP1, PBP3 and PBP4 and the bi-functional transpeptidase/transglycosylase PBP2 responsible for synthesizing the peptidoglycan sacculus) to midcell was recently shown to be driven by the putative lipid II flippase MurJ (Monteiro *et al.*, 2018). The current models of the *S. aureus* cell cycle suggest an initial, gradual increase in cell volume by slight elongation, followed by recruitment of MurJ and PBPs to the septum, which drives cross-wall synthesis and cell constriction. To split the daughter cells, hydrolases perforate the cell wall before the actual splitting/popping of daughter cells occurs on a timescale of milliseconds (Monteiro *et al.*, 2015, Zhou *et al.*, 2015, Lund *et al.*, 2018, Monteiro *et al.*, 2018).

During each cell cycle, peptidoglycan synthesis and cell division need to be coordinated with DNA replication and chromosome segregation. This is to ensure correct cell size homeostasis and that the two daughter cells each get one copy of the chromosome in time before the cell splits. Misregulation would result in daughter cells of variable sizes without DNA or guillotining of the chromosome by the septal cross-wall. How the timing and localization of Z-ring assembly and cell wall synthesis are regulated in *S. aureus*, given the geometry of cell division in three consecutive perpendicular planes, is still an unanswered question. One protein involved in this coordination is probably Noc (nucleoid occlusion protein) which both controls DNA replication (Pang *et al.*, 2017) and inhibits Z-ring formation across the nucleoid (Veiga *et al.*, 2011). Recent results also predict the protein DivIVA to have an important role in linking chromosome segregation with cell division (Bottomley *et al.*, 2017).

The important human pathogen *S. pneumoniae* is an ovococcal bacterium, in which both septal (division) and peripheral (elongation) cell wall synthesis occur in the mid-cell area (Ducret & Grangeasse, 2017). In these cells, positioning of the Z-ring at mid-cell has been shown to depend on several factors, including the chromosomal origin of replication (van Raaphorst *et al.*, 2017) and the peptidoglycan binding protein MapZ (Fleurie *et al.*, 2014a, Holeckova *et al.*, 2014). Most likely, the septal and peripheral cell wall growth in pneumococcal cells are mediated by separate protein machineries, whose actions are tuned by different regulatory proteins such as StkP, MreCD, GpsB, DivIVA or EloR (Ducret & Grangeasse, 2017, Fleurie *et al.*, 2014b, Rued *et al.*, 2017, Straume *et al.*, 2017, Stamsås *et al.*, 2017, Beilharz *et al.*, 2012, Zheng *et al.*, 2017). Another protein involved in regulation of cell wall synthesis in pneumococci, named CozE (for coordinator of zonal elongation, SPD_0768 in strain D39 and Spr0777 in strain R6), was recently identified (Fenton *et al.*, 2016, Straume *et al.*, 2017). CozE, a multi-transmembrane spanning protein, was found to be essential for normal growth, however, its essentiality was abolished in the absence of the bifunctional penicillin-binding protein PBP1a or the cell wall elongation proteins MreC and MreD (Fenton *et al.*, 2016). In protein-protein interaction assays, CozE was found to be associated with the same proteins (PBP1a, MreC, MreD) as well as DivIVA and PBP2b (Fenton *et al.*, 2016, Straume *et al.*, 2017). CozE was thus proposed to be a key regulator of cell elongation in *S. pneumoniae* by positioning PBP1a via interactions with MreC and MreD (Fenton *et al.*, 2016, Ducret & Grangeasse, 2017). CozE proteins are widespread among different bacteria (Fenton *et al.*, 2016). Here we studied the two homologs of CozE in spherical *S. aureus* cells. We show that the CozE proteins are involved in coordinating cell division in *S. aureus* and that this function is conserved also in *S. pneumoniae*.

As a means to study the functionality of essential genes, we also develop a CRISPR interference (CRISPRi) system for *S. aureus*. With CRISPRi, the CRISPR/Cas9-system is harnessed to knock down gene expression of any gene of interest (Bikard *et al.*, 2013, Qi *et al.*, 2013). Transcriptional knockdown is achieved by two components: a catalytically inactive Cas9 protein (dCas9) and a single guide RNA (sgRNA). Unlike the Cas9 nuclease, dCas9 does not cleave DNA, but the DNA-binding capability is still intact. A single guide RNA (sgRNA), containing a gene-specific base-pairing region and a structured region for interaction with dCas9, is designed to target the gene of interest. Upon co-expression, the dCas9-sgRNA complex will bind DNA and serve as a transcriptional roadblock for the RNA polymerase, thereby downregulating transcription.

## Results

### *The two CozE homologs of* S. aureus

The protein CozE (for coordinator of zonal elongation) was recently identified as an essential cell division protein in oval shaped pneumococcal cells, where it has been shown to be involved in regulation of proper cell elongation (Fenton *et al.*, 2016, Straume *et al.*, 2017). Spherical *S. aureus* does not elongate to the same extent as *S. pneumoniae* and other rod-or oval-shaped bacteria, although a short elongation phase has been observed during the cell cycle (Pinho *et al.*, 2013, Monteiro *et al.*, 2015). Nevertheless, homology searches showed that *S. aureus* encodes two proteins homologous to CozE, and we therefore set out to unravel the function of CozE in these spherical cells. Sequence comparison of the pneumococcal CozE (hereafter CozE^Spn^) with the two CozE-homologs of *S. aureus* SH1000, SAOUHSC_00948 (hereafter CozEa) and SAOUHSC_01358 (hereafter CozEb), shows that they are 31 % and 30 % identical to CozE^Spn^, respectively (Fig. S1). When compared with each other, CozEa and CozEb are 30 % identical. Topology predictions suggest that these proteins have 8 or 9 transmembrane segments (Fig. S1). The *cozEa* gene is predicted to be monocistronic, while *cozEb* is located as the last open reading frame on a three-gene operon which also encodes a transcription antiterminator (*glcT*) and a small, putative membrane spanning protein (SAOUCHSC_01357) (Fig. 1A).

**Fig. 1.**
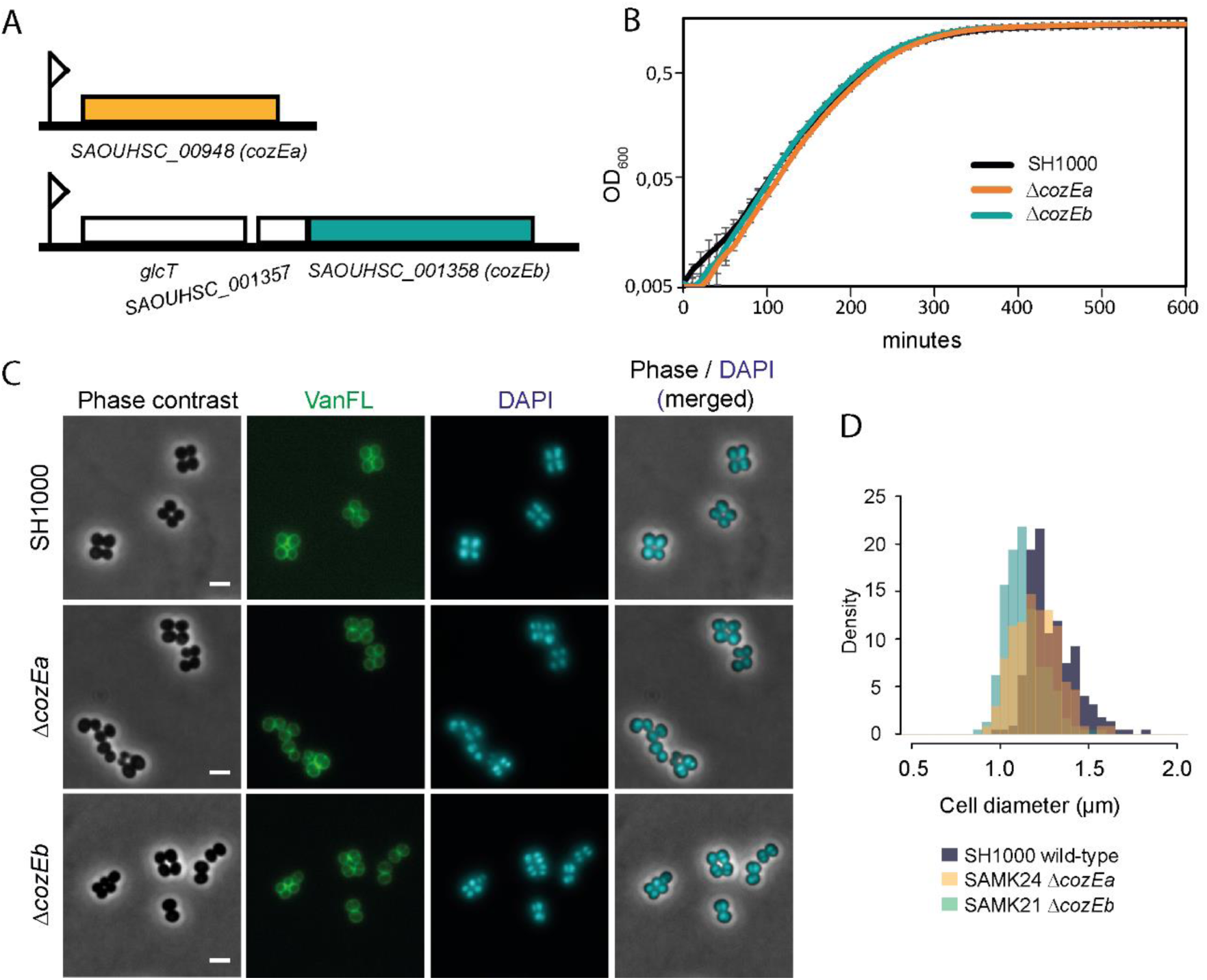
*cozEa* and *cozEb* of *S. aureus*. A.Genetic organization of the *cozEa* (SAOUHSC_00948) and *cozEb* (SAOUHSC_01358) genetic loci. B.Growth curves SH1000 wild-type, SAMK24 (Δ*cozEa*) and SAMK21 (Δ*cozEb*) in BHI medium at 37°C. C.Micrographs of SH1000, SAMK21 and SAMK24. Phase contrast (PC) images and staining with fluorescent vancomycin (VanFL) and DAPI are shown as well as an overlay of the two latter. The scale bars are 2 μm D.Histogram of the cell diameters of SH1000, SAMK24 and SAMK21 (>250 cells analyzed per sample) as measured using MicrobeJ (Ducret *et al.*, 2016). Both SAMK21 and SAMK24 were significantly smaller than SH1000 (P < 0.05, Kolmogorov-Smirnov test).

Using the temperature sensitive vector pMAD (Arnaud *et al.*, 2004), *cozEa* and *cozEb* were deleted individually in *S. aureus* SH1000 by allelic exchange with a spectinomycin resistance cassette. The deletion mutants SAMK24 (Δ*cozEa*∷*spc*) and SAMK21 (Δ*cozEb*∷*spc*) did not exhibit any growth defect compared to wild-type (Fig. 1B). Analysis of cell sizes showed that the cell diameter of both mutants, on average, are slightly smaller compared to the wild-type (Fig. 1C and D). No obvious differences in cell wall labelling (using fluorescent vancomycin, VanFL) or nucleoid staining patterns (using DAPI) were observed between the mutants and wild-type (Fig. 1C).

In order to see whether the two mutant strains, SAMK21 and SAMK24, had acquired any suppressor mutations elsewhere in the genome, we resequenced their genomes and compared it to the SH1000 wild-type genome. SAMK24 did not contain any additional mutations. In SAMK21, a single conservative SNP was found in the gene *thiI* (SAOUHSC_01824) encoding a probable tRNA sulphurtransferase. This SNP (A970T) resulted in a conservative substitution of isoleucine with a phenylalanine (I324F). Our later experiments (see below) show that this mutation is not important for the functionality of *cozEb* (or *cozEa*) and we therefore conclude that neither CozEa nor CozEb are essential for normal growth and cell division in *S. aureus* SH1000.

Single deletions of *cozEa* or *cozEb* both cause a small reduction in cell size. To investigate the effects of a double deletion, another pMAD deletion vector (pMAD-*cozEa*∷*cam*) was constructed to delete *cozEa* in the Δ*cozEb*∷*spc* background. However, despite multiple attempts, we were unable to obtain the double deletion strain. This suggests that *cozEa* and *cozEb* may have complementary and essential functions.

### *Construction of a two-plasmid CRISPR interference system for* S. aureus

Since double deletions of *cozEa* and *cozEb* could not be obtained, we instead wanted to study the phenotypes of the cells when *cozEa* or *cozEb* gene expression was knocked down in Δ*cozEb* or Δ*cozEa* background, respectively. We therefore constructed a CRISPR/dCas9 knockdown system to allow inducible depletion of essential genes. The CRISPR interference systems developed for *S. pneumoniae* and *Bacillus subtilis* (Liu *et al.*, 2017, Peters *et al.*, 2016) were used as models. A *dcas9* gene, encoding a catalytically inactive Cas9, was cloned downstream of an IPTG-inducible promoter in the low-copy number plasmid pLOW (pSK41 minireplicon, Fig. 2A) (Liew *et al.*, 2011). A single guide RNA (sgRNA) construct, consisting of a 20 nt base-pairing region and a Cas9-handle region, was inserted downstream of a synthetic, constitutive promoter in the plasmid pCG248 (replicon T181, Fig. 2A) (Helle *et al.*, 2011). Targeting of the gene of interest is accomplished by replacing the 20 nt sequence using inverse PCR as described in the Methods section. Notably, multi-sgRNA plasmids can be constructed by using the BglII and BamHI restriction sites located up- and downstream of the sgRNA construct, as outlined in Fig. S2. A schematic view of the resulting two-plasmid CRISPRi system is shown in Fig. 2A.

**Fig. 2.**
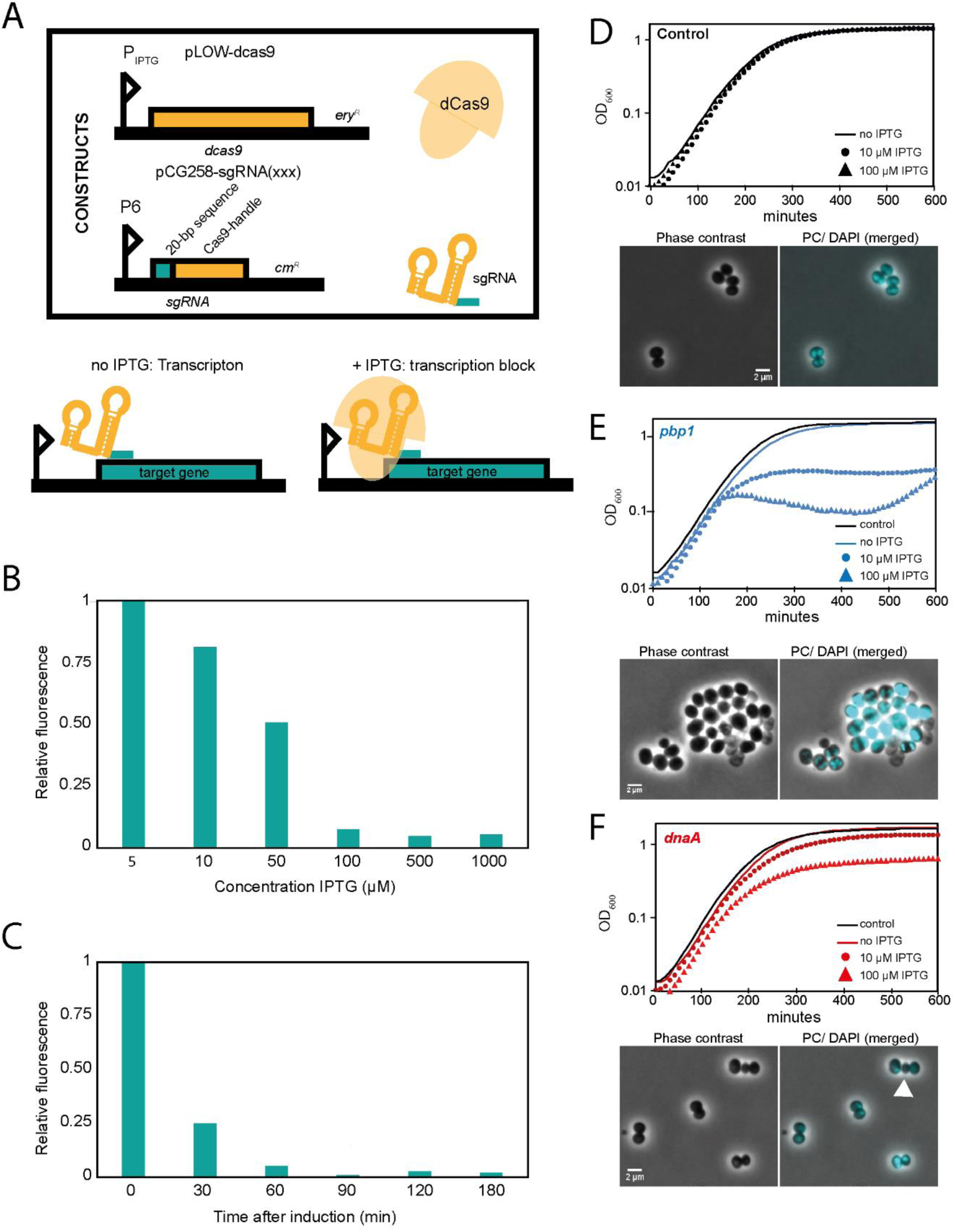
Two-plasmid CRISPR interference system for *S. aureus*. A.Schematic representation of the plasmids carrying dCas9 and sgRNA. The sgRNA is constitutively expressed, while the level of dCas9 is controlled by the inducible P_*lac*_ promoter. Upon addition of IPTG, dCas9 will be expressed and the dCas9-sgRNA-DNA complex formation will lead to transcription block and knockdown of the target gene. B-C. Knockdown of GFP expression in a strain constitutively expressing *m(sf)gfp* (SAMK56). B.Fluorescence after induction with various IPTG concentrations. The fluorescence values are given relative to the fluorescence of a non-depleted strain. The experiment was repeated twice with similar results. C.The temporal dynamics of GFP depletion after addition of 100 μM IPTG. The fluorescence at the time of IPTG addition was set to 1, and measured at different time points. The experiment was repeated twice with similar results. D-F. Growth and phenotypic characterization of cells with depletion using CRISPRi. Growth curves in BHI medium at 37°C and micrographs are shown. The cultures were diluted to OD_600_ ∼ 0.01 prior to growth analysis. The scale bars are 2 μm. D.Control cells carrying non-targeting sgRNA. E.Depletion of *pbp1. pbp1* depleted cells were significantly larger than wild-type cells (1.78 ± 0.38 μm, n = 126 for the *pbp1* depletion versus 1.41 ± 0.34 μm, n = 250 for the control, P < 0.05, Kolmogorov-Smirnov test). F.Depletion of *dnaA*, resulted in formation of anucleate cells (10.2 %, n = 234). The arrowhead points to an example of an anucleate cell.

To quantify the efficiency of our CRISPRi system, we created an RN4220-derivative strain with constitutive expression of a monomeric superfolder GFP (m(sf)GFP), SAMK56, and designed an sgRNA targeting the *m(sf)gfp* gene. As shown in Fig. 2B, GFP expression could be titrated by increasing the IPTG concentrations. Maximum depletion was obtained with ≥100 μM IPTG. To investigate how quick GFP expression was switched off after IPTG induction, SAMK56 was induced with 100 μM IPTG and samples were taken every 30^th^ min for 3 hrs. The GFP fluorescence levels (Fig. 2C) decreased rapidly (signal reduced by ca. 90% within 60 min), suggesting that expression was switched off almost immediately. Specific transcriptional knockdown was also demonstrated using qPCR (Fig. S3, see below for details). Note that for some of our later experiments, we observed a faster depletion of cell division proteins when increasing the IPTG concentration to 400 μM. Furthermore, as a proof of the functionality of the CRISPRi system in targeting essential cell cycle genes, we created sgRNAs targeting the DNA replication initiator *dnaA* (encoded on an operon with *dnaN*) and *pbp1* (monocistronic) encoding a penicillin-binding protein. The CRISPRi strains were analyzed by growth assays and microscopy (Fig. 2D-F), and the observed phenotypes were as expected, confirming the suitability of the CRISPRi system to study the function of essential genes; compared to the control strain (Fig. 2D), the *pbp1* depletion resulted in clustered, larger cells with aberrant morphologies (Fig. 2E) (Pereira *et al.*, 2007, Pereira *et al.*, 2009) while *dnaA* depletion resulted in anucleate cells with variable sizes and nucleoid morphologies (Fig. 2F).

### *CozEa and CozEb have overlapping functions and are important for proper cell cycle progression in* S. aureus

We made sgRNA constructs targeting *cozEa* and *cozEb*, and depleted expression of *cozEa* in the Δ*cozEb* background and vice versa. Note that *cozEa* is monocistronic, while *cozEb* is located as the last gene in the operon, and the knockdown will therefore have minimal polar effects (Peters *et al.*, 2016, Liu *et al.*, 2017). No growth reduction was observed upon knockdown of the individual genes in wild-type background (as expected from the deletion mutants) (Fig. 3A and B). We performed RT-qPCR on these CRISPRi-strains and verified that transcription of *cozEa* and *cozEb* was specifically knocked down (Fig. S3). After diluting the cells to OD_600_ of 0.05 with and without 150 μM IPTG and culturing for 2 hours, the expression of *cozEa* and *cozEb* was reduced 23-fold and 13-fold in the respective strains when comparing induced and non-induced conditions.

**Fig. 3.**
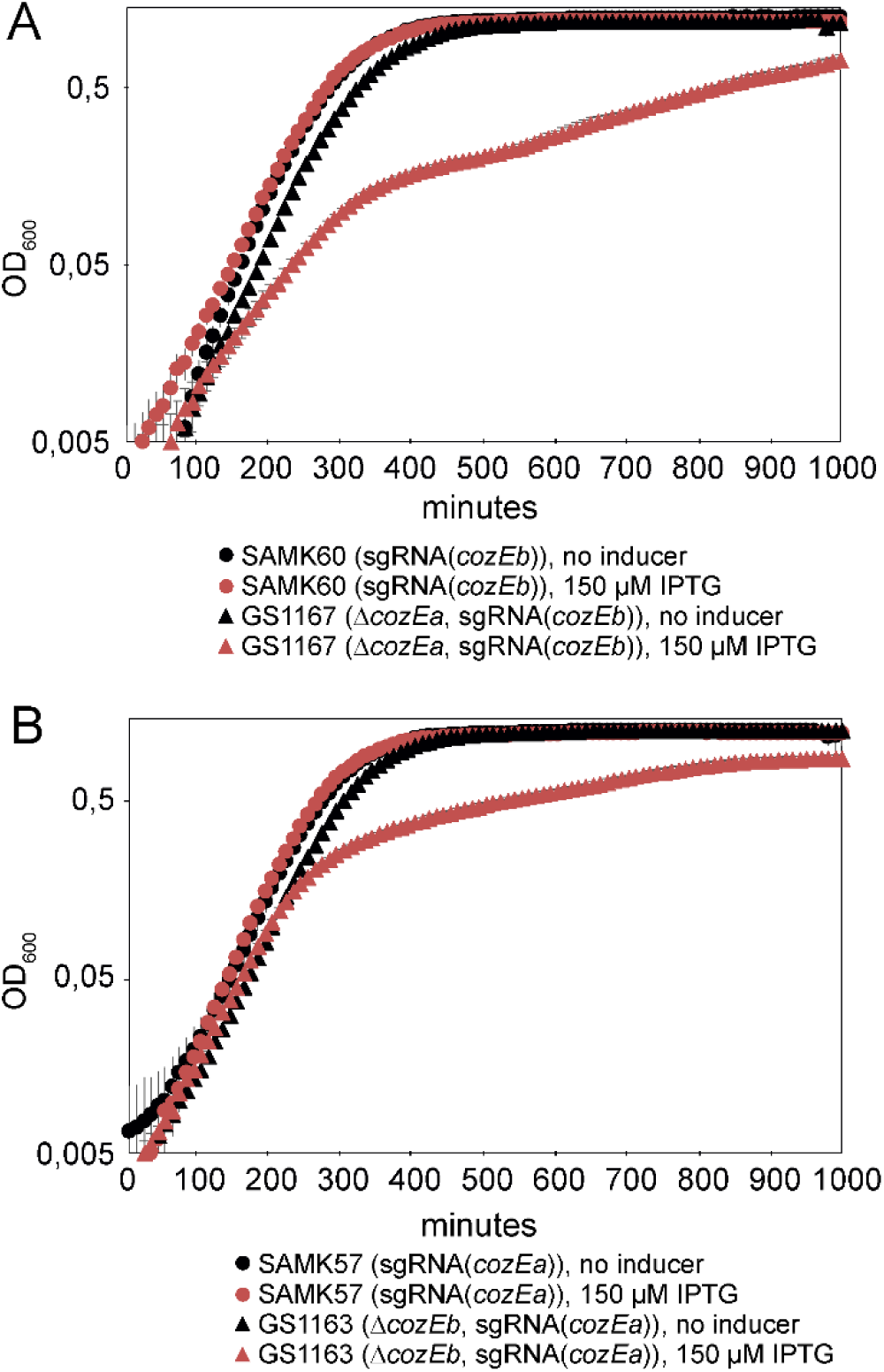
Functional redundancy of *cozEa* and *cozEb*. A.Strains carrying sgRNA targeting *cozEb* in wild-type background (circles, strain SAMK57) compared to the Δ*cozEa* background (triangles, strain GS1167). Growth curves with (red) or without (black) induction of dCas9 expression with 150 μM IPTG are shown. B.Strains carrying sgRNA targeting *cozEa* in wild-type background (circles, strain SAMK60) compared to the Δ*cozEb* strain (triangles, strain GS1163). Growth curves with (red) or without (black) induction of dCas9 expression with 150 μM IPTG are shown.

While no effect on growth was observed by depleting *cozEa* or *cozEb* expression in wild-type background, knockdown of the other gene in the respective deletion backgrounds caused dramatic reduction in growth (Fig. 3A and B). The initial doubling time after CRISPRi-induction is more affected in GS1167 (Δ*cozEa*, sgRNA(*cozEb*), t_d_ ^induced^ = 52 min and t_d_ ^non-induced^ = 34 min) than in GS1163 (Δ*cozEb*, sgRNA(*cozEa*), t_d_^induced^= 38 min and t_d_^non-induced^= 35 min). However, after approximately 300 min of dCas9-induction, the growth is dramatically reduced for both GS1167 and GS1163 (Fig. 3). From this we conclude that *cozEa* and *cozEb* have overlapping functions in *S. aureus*.

The phenotypes of the GS1167 (Δ*cozEa*∷*spc*, depleted *cozEb*) and GS1163 (Δ*cozEb*∷*spc*, depleted *cozEa*) strains were then further investigated by microscopy. Phase contrast micrographs revealed severely perturbed cell morphologies when the CRISPRi system is induced, displaying both variable cell shapes and sizes as well as increased clustering of cells (Fig. 4A). Measurements of the cell diameter of CRISPRi-induced GS1167 and GS1163 cells show that they have a very wide distribution compared to the wild type (Fig. 4B). We also made a double sgRNA strain allowing knockdown of both *cozEa* and *cozEb* simultaneously with the CRISPRi system (strain SAMK75), and as expected this strain displayed similar phenotype as the GS1167 and GS1163 strains (Fig. S4).

**Fig. 4.**
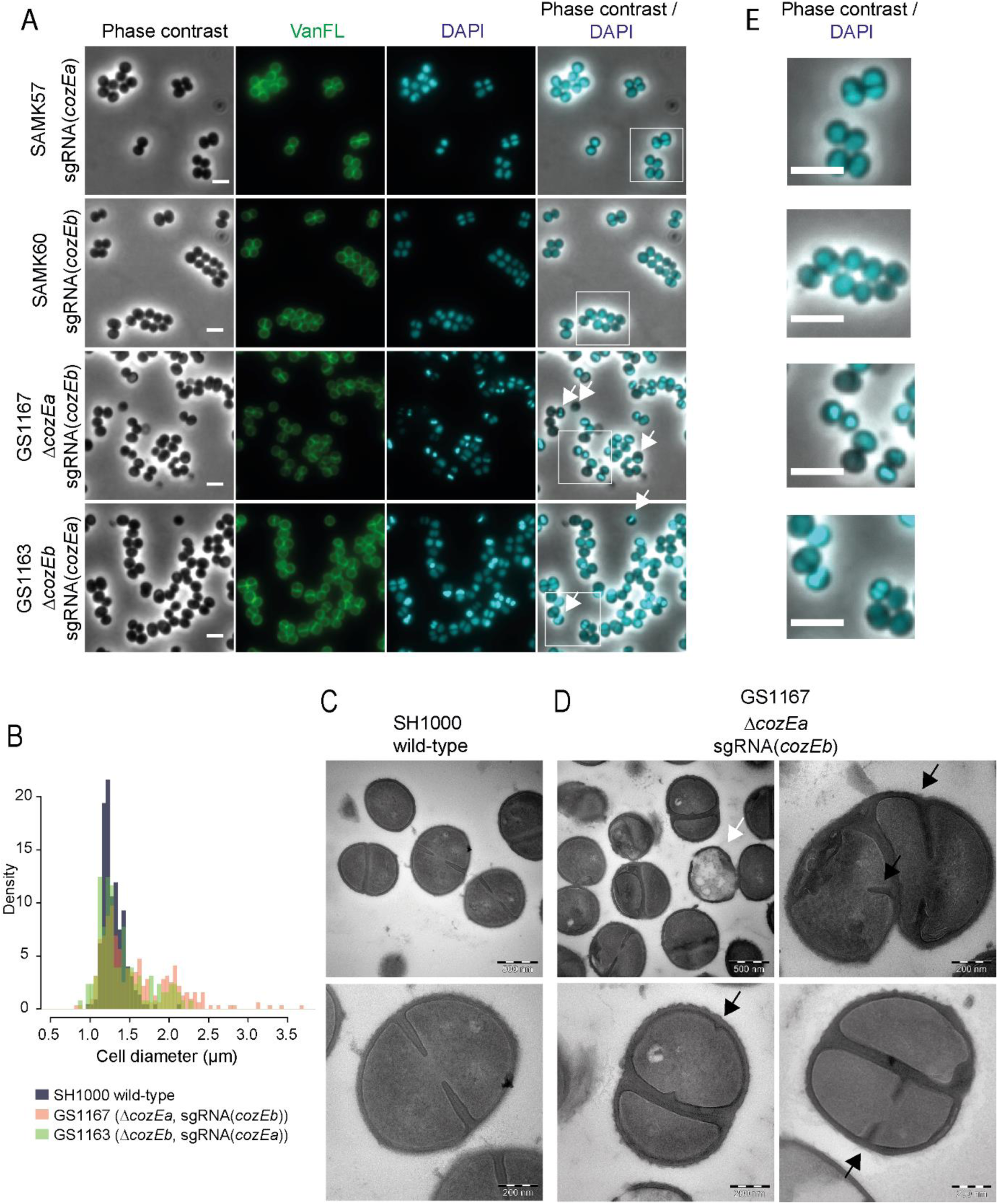
CozEa/CozEb phenotypes in *S. aureus* SH1000. A.Phase contrast micrographs are shown to the left. The cells were also stained with fluorescent vancomycin and DAPI to visualize cell wall and the nucleoid, respectively. Overlay of DAPI and phase contrast images are shown. White arrows point to cell with aberrant nucleoids. The scale bars are 2 μm. The white squares indicate the area magnified in Fig. 4E. B.Histogram of cell diameter distribution for wild-type cells (grey) and Δ*cozEa*∷*spc* with depleted *cozEb* (GS1167, orange) cells and Δ*cozEb*∷*spc* with depleted *cozEa* (GS1163, green) induced with 150 μM IPTG. Both GS1167 and GS1163 are different from wild-type (P < 0.05, Kolmogorov-Smirnov test), with high proportion of the cells with diameters larger than 1.5 μm (7.9 % for wild-type compared to 44.2% for GS1167 and 27.9 % for GS1163, n >150 for all strains). C-D. Transmission electron micrographs of wild-type cells (C, SH1000) and Δ*cozEa*∷*spc* with depleted *cozEb* (D, GS1167) cells. The white arrow points to a lysed cell. Black arrows point to septum initiation in GS1167 cells. Two different magnifications are shown, as indicated by the scale bars. E. Magnified insets from Fig. 4A with overlays of DAPI and phase contrast images, demonstrating the variation in nucleoid staining between the strains.

Perturbed morphologies in cells depleted of both *cozEa* and *cozEb* prompted us to further analyze cell division placement by transmission electron microscopy (TEM) (Fig. 4C - D and S4 Fig.). In GS1167 cells (Δ*cozEa*∷*spc* with depleted *cozEb*) depleted for 4 hours, cells could initiate septum formation in only one of the daughter cells prior to cell splitting (Fig. 4D, lower panels) and non-perpendicular septum formation resulting in misshaped cells was also observed (Fig. 4D, top right panel). Spatial and temporal coordination of cell division thus seem compromised in cells lacking CozEa and CozEb. Empty, lysed cells were also observed (Fig. 4D). Furthermore, the cell wall also appeared to be thicker in the mutant cells, and comparison of septum thickness based on the TEM images show that the GS1167 on average has thicker septal cross-wall compared to wild-type cells (Fig. S6). In mildly depleted cells (depletion for 1 hour), the phenotype is less severe, however, uncoordinated initiation of septum formation and aberrant septa were also observed here (Fig. S5). Electron micrographs of the individual deletions showed that Δ*cozEa* mutants display both lysed cells (Fig. S5) and have thicker septa than wild-type (Fig. S6). These phenotypes were not observed in Δ*cozEb* cells (Fig. S5 and S6).

Notably, nucleoid staining of the cells depleted of both *cozEa* and *cozEb* using DAPI was also abnormal, displaying non-homogeneous staining patterns. A large fraction of the cells appeared to have high intensity or highly condensed DAPI signals (47.1 % for GS1167, 22.9 % for GS1163, n > 250) compared to the wild type, and some cells were also anucleate under these conditions (4.1 % for GS1167 and 2.0 % for GS1163, n > 250) (Fig. 4A and zoomed images in Fig. 4E). The chromosome biology of the cells thus also seems to be perturbed when CozEa and CozEb are lacking.

### CozEa and CozEb do not affect cell wall composition, but interact with key cell division proteins

TEM images showed that the septal cell wall appeared different between wild-type and the *cozE*-deficient cells; coordination of cell wall synthesis seems to be compromised (Fig. 4) and the septal cell wall is thicker in cells lacking *cozEa* (Fig. S5.). In *S. pneumoniae*, CozE^Spn^ has been shown to interact with the bi-functional penicillin binding protein PBP1a (Fenton *et al.*, 2016). To get insight into whether CozEa and CozEb could influence cell wall synthesis, we first investigated whether these proteins could interact with any of the four PBPs of *S. aureus* (PBP1, PBP2, PBP3 and PBP4) using bacterial two-hybrid assays (see Material and Methods for detailed description) (Karimova *et al.*, 2005). While CozEa and CozEb both self-interacted and interacted with each other, no interaction was found with any of the PBPs of *S. aureus* (Fig. 5A, Fig. S7). We also tested the methicillin-resistant PBP2a (MecA) from *S. aureus* COL, but we could not find any interactions with the CozE proteins (Fig. 5A). Next, we analyzed the muropeptide composition of peptidoglycan derived from strain the GS1167 (Δ*cozEa* with depleted *cozEb*), to see whether the cell wall architecture was altered in this mutant. However, the muropeptide composition of GS1167 was similar to the wild-type (Fig. 5B). This suggests that CozEa and CozEb affect positioning and timing of cell division and cross-wall synthesis, but that the cell wall synthesis pathway is unaltered.

**Fig. 5.**
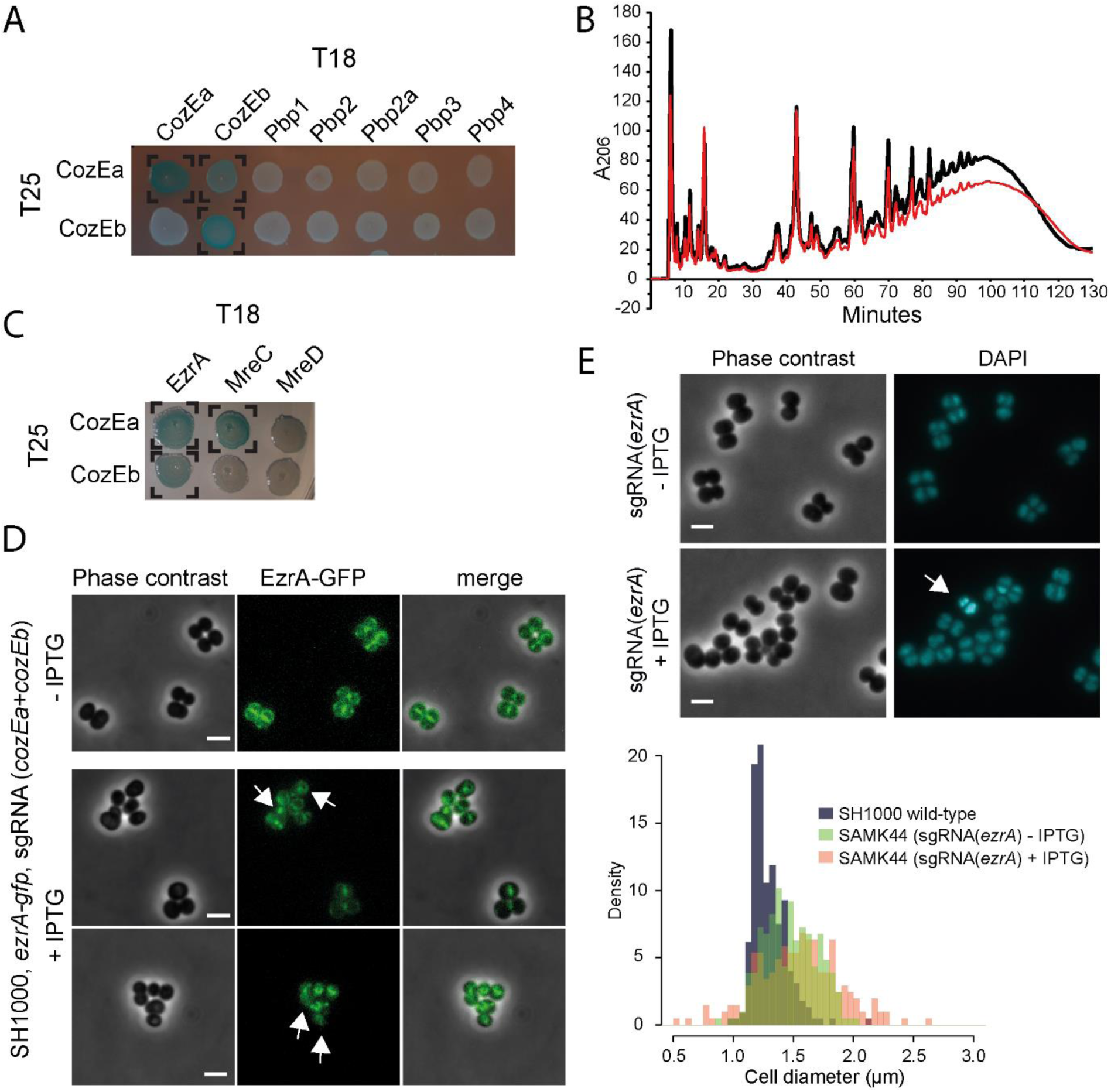
CozEa and CozEb interact with EzrA, but do not alter cell wall synthesis. A. Bacterial two-hybrid analyses of interactions between CozEa and CozEb fused to the T25 domain with proteins fused to the T18. Positive interactions are observed as blue colonies and marked with brackets. See Fig. S7 for control experiments. All interaction results were repeated at least five times. B. Cell wall muropeptide composition of SAMK15 control cells (black) and GS1167 depletion cells (red) induced with 150 mM IPTG for 4 hours as analyzed with UHPLC. See Fig. S7 for control experiments. C. Bacterial two-hybrid analyses of interactions between CozEa and CozEb fused to the T25 domain with EzrA, MreC and MreD fused to the T18 domain. Positive interactions are observed as blue colonies and marked with brackets. See Fig. S7 for control experiments. All interaction results were repeated at least five times. D. Localization of EzrA-GFP without and with induction of CozEa/CozEb-depletion. The upper panel shows uninduced cells and two lower panels show representative cells after induction of CRISPRi with 400 μM IPTG. The arrows point to cells with obvious mislocalization of EzrA-GFP. The scale bars are 2 μm. E. Phenotype of *ezrA* knockdown. Phase contrast micrographs and DAPI signal are shown for SAMK44 (CRISPRi targeting *ezrA*) with or without induction with 300 μM IPTG. The arrows point to cells with aberrant nucleoid staining. The scale bars are 2 μm. The lower panel shows cell diameter histograms of wild-type SH1000 cells as well as induced and non-induced SAMK44 cells. Both induced and non-induced cells are significantly larger than wild-type cells (Kolmogorov-Smirnov test, P < 0.05), with high proportion of the cells with diameters larger than 1.5 μm (7.9 % for wild-type compared to 43.9% for non-induced and 55.1 % for induced, n >100 for all strains).

We further analyzed whether CozEa and CozEb could interact with a selection of other key cell cycle proteins using bacterial two-hybrid assays (Fig. 5C, S1 Table, Fig. S7). CozE^Spn^ has been shown to interact with MreC^Spn^, MreD^Spn^ and DivIVA^Spn^ (Straume *et al.*, 2017, Fenton *et al.*, 2016). We detected an interaction between CozEa and MreC, however, this was not the case for CozEb. No interactions with MreD or DivIVA were observed for any of the combinations (Fig. 5C, S1 Table). The only protein we could identify that interacted with both CozEa and CozEb was the early cell division protein EzrA. Among the other proteins tested, we also found that CozEa and GpsB interacted, however, not CozEb and GpsB. A full overview of all tested bacterial two-hybrid interactions are given in S1 Table.

The positive two-hybrid interactions suggest that CozEa and CozEb may mediate cell division control via interactions with EzrA. CozEa and CozEb both displayed a membrane localization in *S. aureus* SH1000, with no apparent enrichment in the septal region (Fig. S9). Notably, however, depletion of *cozEa* and *cozEb* expression in cells expressing a chromosomal *ezrA-gfp* fusion (Lund *et al.*, 2018), demonstrate that EzrA-GFP is mislocalized under these conditions (Fig. 5D). Furthermore, knockdown of *ezrA* using the CRISPRi system leads to similar phenotypes as cells lacking CozEa and CozEb (Fig. 5E) with variable cell sizes and nucleoid staining. This is fully in line with previous results of *ezrA* deletions and knockdown mutants in *S. aureus* (Steele *et al.*, 2011, Jorge *et al.*, 2011). Note that the cell size effect is observed also, but to a lesser extent, when no IPTG is added, reflecting leaky expression from the P_*lac*_ promoter. Abnormal DAPI staining pattern was also observed in the cells after IPTG induction, although this phenotype appear to be less pronounced in the *ezrA* knockdown cells (5.5 % of cells, n = 200) compared to cells depleted of CozEa and CozEb. It should also be noted that the growth rate was not severely affected upon induction of *ezrA* knockdown (Fig. S6), suggesting that *ezrA* is not essential for normal growth under these conditions (Bottomley *et al.*, 2014).

### *The division ring is mislocalized in* S. pneumoniae *cells depleted of CozE*^*Spn*^

The results above demonstrate that CozEa and CozEb play functionally overlapping roles in controlling cell division in *S. aureus*, and both genes can be deleted individually. As mentioned above, a single protein CozE^Spn^ (SPD_0768 in strain D39 and Spr0777 in strain R6), is shown to be essential for growth and proper cell morphology in *S. pneumoniae* (Fenton *et al.*, 2016, Straume *et al.*, 2017). To investigate whether the EzrA-interactions detected here were specific for *S. aureus* or also relevant in *S. pneumoniae*, we used bacterial two-hybrid assays to test the interaction between CozE^Spn^ and EzrA^Spn^. Just like the staphylococcal proteins, a strong interaction was found between the corresponding pneumococcal proteins (Fig. 6A). Strikingly, while EzrA localized to midcell in wild-type *S. pneumoniae* (Fig. 6B and C), the protein is clearly mislocalized in cells where *cozE*^*Spn*^ was depleted (Fig. 6D).

**Fig. 6.**
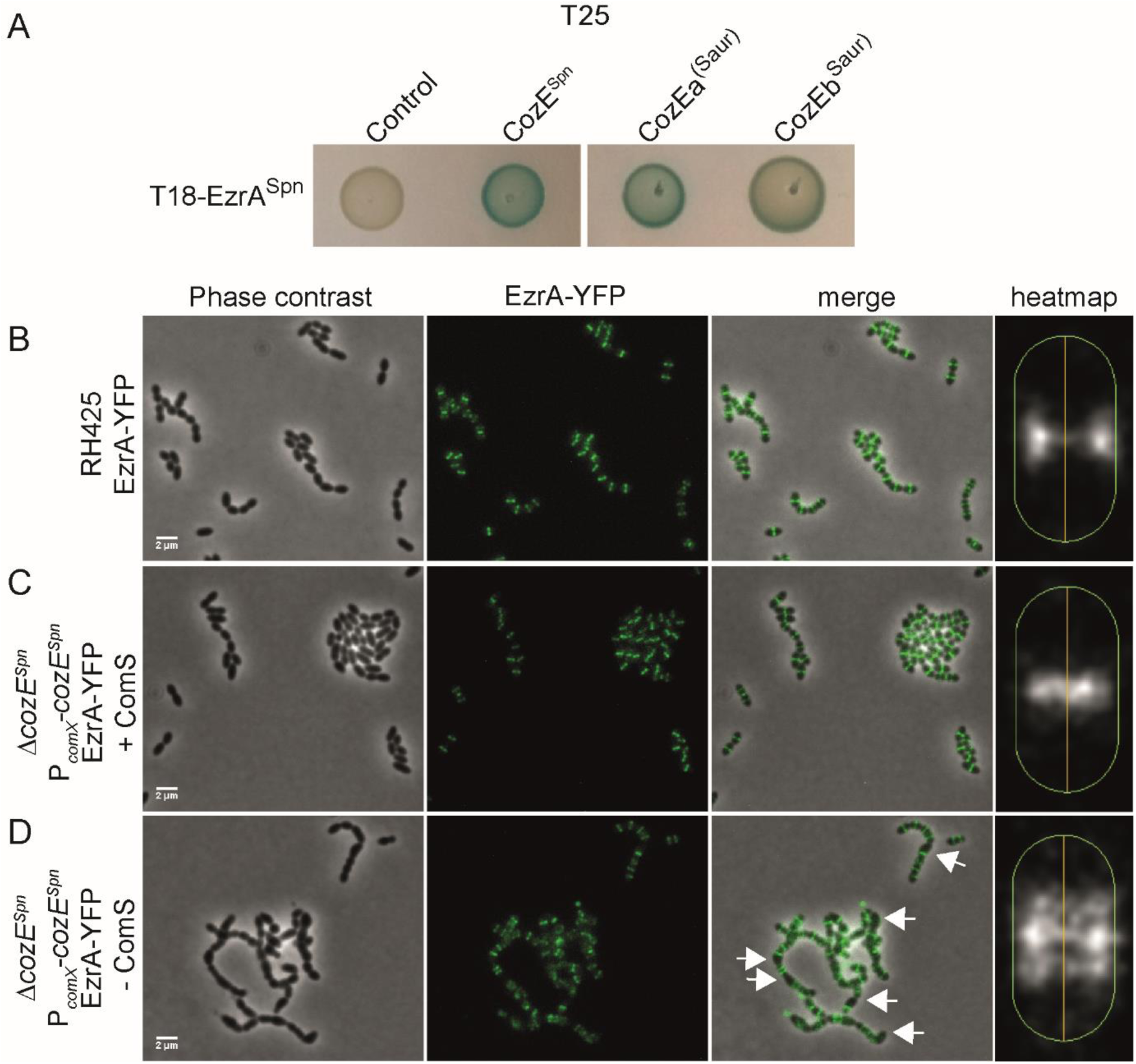
CozE^Spn^ controls division ring formation in *S. pneumoniae*. A. Bacterial two-hybrid assay showing interactions between EzrA^Spn^ and different CozE-proteins. B-D. Localization of EzrA-YFP in *S. pneumoniae*. Phase contrast and fluorescence are shown individually and merged. The localization of EzrA-YFP in the cells are also shown as heatmaps, as generated using MicrobeJ. The heatmaps represent the localizations in >650 cells for each strain. EzrA-YFP localization was analyzed in RH425 wild-type (B) and a strain where the native *cozE*^*Spn*^ is deleted and instead expressed from the ComS-inducible promoter P_*comX*_ (Berg *et al.*, 2011). The *cozE*^*Spn*^-depletion strain (Δ*cozE*^*Spn*^ and P_comX_-*cozE*^*Spn*^) was grown with (C) or without (D) inducer peptide ComS. The arrows point to examples of cells with mislocalized EzrA-YFP. The scale bars are 2 μm.

S. aureus cozEa *and* cozEb *can complement the* ΔcozE ^Spn^ *phenotype of* S. pneumoniae In order to gain further insight into functional conservation of CozE proteins between *S. aureus* and *S. pneumoniae*, we tested whether CozEa or CozEb could functionally complement the essential CozE^Spn^ in *S. pneumoniae*. We created pneumococcal strains in which *cozEa* and *cozEb* were chromosomally integrated downstream of the ComRS-inducible promoter, P_*comX*_. Induction of P_*comX*_ is achieved by addition of the peptide ComS to the growth medium; ComS is internalized where it activates the P_*comX*_-binding transcriptional activator ComR (Berg *et al.*, 2011). Next, we attempted to delete the native *cozE*^*Spn*^ by allelic exchange with the Janus cassette (Sung *et al.*, 2001), with and without presence of the inducer ComS. A functional complementation with CozEa or CozEb in the pneumococcus would allow deletion of the *cozE*^Spn^ gene. Indeed, upon induction of *cozEa* or *cozEb* expression with 2 μM ComS, the native *cozE*^*Spn*^ could readily be deleted (Table 1). It should be noted that the CozEa and CozEb probably have a reduced functionality compared to CozE^Spn^, as higher inducer concentrations were required to obtain correct transformants for the non-native CozE-proteins (Table 1). Additionally, CozEa seemed to function better than CozEb, since the number of transformants were higher for the former. Microscopy of the resulting strains further confirmed that the typical *cozE*^*Spn*^-depletion phenotype in pneumococci, characterized by extensive chaining and slight rounding of cells (seen by reduced length and aspect ratio closer to 1) (Straume *et al.*, 2017), could be complemented by both CozEa or CozEb (Fig. S10).

**Table 1.** Complementation of *ΔcozE*^*Spn*^ in *S. pneumoniae* with CozEa and CozEb.

| **Complementation** | **Colonies/ml/ng<sup>a</sup>** |  |  |
| --- | --- | --- | --- |
|  | **0 $\mu\text{M}$ ComS** | **0.2 $\mu\text{M}$ ComS** | **2 $\mu\text{M}$ ComS** |
| $P_{\text{comX}}\text{-}\text{cozE}^{Spn}$ | 595 (0/8)<sup>b</sup> | >2000 (8/8) | >2000 (8/8) |
| $P_{\text{comX}}\text{-}\text{cozEa}$ | 65 (0/8) | 580 (3/8) | >2000 (8/8) |
| $P_{\text{comX}}\text{-}\text{cozEb}$ | 71 (0/8) | 77 (0/8) | 45 (6/8) |

<sup>a</sup> number of colonies on the plate after 16 hours (per 1 ml transformation mix per ng DNA) when the respective strains were transformed with the DNA fragment $\Delta\text{cozE}^{Spn}\text{:P-}\text{rpsL-kan}$ (Janus cassette). Transformants were selected with kanamycin. Eight colonies for each transformation were checked by PCR, and the number of true $\Delta\text{cozE}^{Spn}$ verified by PCR per 8 colonies are indicated in brackets.
<sup>b</sup> When transforming with the $\Delta\text{cozE}^{Spn}\text{:P-}\text{rpsL-kan}$ cassette, small sized colonies are observed also without complementation. However, these are not true $\Delta\text{cozE}^{Spn}$ deletions when checked by PCR. In these colonies, $\text{cozE}^{Spn}$ has moved to another chromosomal location (data not shown) and the strain has probably acquired suppressor mutations as previously observed.

Finally, to get insights into how the staphylococcal proteins CozEa and CozEb could complement CozE^Spn^, we analyzed by bacterial two-hybrid assays whether CozEa and CozEb could still interact with EzrA^Spn^ (Fig. 6A). Both CozEa and CozEb interact with EzrA^Spn^ in this assay. Thus, conservation of the interaction with EzrA could thus explain why CozEa and CozEb were functional in *S. pneumoniae*.

## Discussion

The membrane protein CozE^Spn^ was recently identified as an essential regulator of cell elongation in oval shaped *S. pneumoniae* (Fenton *et al.*, 2016, Straume *et al.*, 2017). CozE proteins are widely conserved and present in the genome of bacteria from different phyla and of different morphologies (Fenton *et al.*, 2016). We here show that the two CozE-homologs of *S. aureus*, which we named CozEa and CozEb, play overlapping roles to control proper cell cycle progression in these spherical cells.

While the deletion of either *cozEa* or *cozEb* has only minor effects, both genes cannot be deleted at the same time. To confirm the synthetic relationship between *cozEa* and *cozEb*, we developed a CRISPRi system for *S. aureus* to allow knockdown of expression of essential genes. Recent reports have already shown the suitability of using CRISPR/dCas9 for knockdown of genes in *S. aureus* (Dong *et al.*, 2017, Zhao *et al.*, 2017). The plasmid-based CRISPR/dCas9-derived system we developed here contains several unique features compared to the published ones (Dong *et al.*, 2017, Zhao *et al.*, 2017): (i) Knockdown is inducible by addition of IPTG, since dCas9 expression is driven by the IPTG-inducible promoter. (ii) The plasmid harbouring the sgRNA construct is relatively small (5.8 kb), thus allowing easy replacement of target sequences by inverse PCR. (iii) Multi-sgRNA plasmids, allowing simultaneous knockdown of several genes, can be constructed by combining existing sgRNA plasmids using BglBrick assembly (Anderson *et al.*, 2010) (Fig. S2).

Using the CRISPRi system, we could construct combined deletion/depletion strains or double-depletion strains to study cells depleted of CozE proteins. Since all the different strains depleted of CozE proteins showed the same phenotypes, we could exclude that the conservative substitution in the gene *thiI* (detected by whole genome resequencing, see results) played any functional role. Low levels of CozEa and CozEb proteins have pleiotropic effects on the staphylococcal cells, including abnormal cell size homeostasis and nucleoid staining, frequent lysis and, most strikingly, the thickened cell wall and compromised timing and positioning of cell division (Fig. 4). Wild-type *S. aureus* cells divide in consecutive, perpendicular planes, i.e. the new septum is formed perpendicular to the previous and splitting of daughter cells (Monteiro *et al.*, 2018) (popping) finishes before the next septum is formed (Pinho *et al.*, 2013, Zhou *et al.*, 2015, Monteiro *et al.*, 2015). However, cells lacking CozEa and CozEb can initiate septum formation asynchronously in only one of the daughter cells before the previous division cycle finishes and non-perpendicular septa were also observed, resulting in elongated cells. This is reminiscent of elongating staphylococcal FtsZ mutant strains (Fig. 4) (Pereira *et al.*, 2016) or staphylococci treated with antibiotics targeting the cell wall or cell division (Lund *et al.*, 2018, Pinho *et al.*, 2000, Sieradzki & Tomasz, 2006).

Despite having misplaced and thicker septa than wild-type, the cell wall composition does not appear to be altered in the CozEa/CozEb-depleted cells and the membrane proteins CozEa or CozEb are not directly interacting with any of the PBPs of *S. aureus*. An interaction between CozEa and MreC was detected, however, CozEa nor CozEb could interact with MreD. Although the CozEa-MreC interaction may be important for directing peptidoglycan synthesis, like in *S. pneumoniae*, it is worth noting MreC and MreD are non-essential in *S. aureus* (Tavares *et al.*, 2015). The significance of the CozEa-MreC interaction thus remains unknown.

CozEa and CozEb might compromise cell division coordination and autolytic splitting by interfering directly with key cell division proteins. The detailed mechanism of action remains to be determined, but we show that CozEa and CozEb could interact with one of the early cell division proteins, namely EzrA, in two-hybrid interaction assays. Notably, the EzrA-GFP localization in *S. aureus* cells lacking CozEa and CozEb was severely perturbed. Interaction with EzrA could thus be a plausible way for CozEa and CozEb to mediate cell division control. EzrA is one of the first proteins binding to the Z-ring in the initiation of cell division. EzrA was identified as a negative regulator of FtsZ formation in *B. subtilis* (Levin *et al.*, 1999), and is thought to be important for the switch between elongation and division growth in *B. subtilis* via protein-protein interactions with penicillin-binding proteins (Claessen *et al.*, 2008). EzrA plays a similar role in *S. pneumoniae* (Rued *et al.*, 2017). In *S. aureus*, EzrA is involved in a large number of protein-protein interactions. Bacterial two-hybrid interactions have been shown between EzrA and FtsZ, DivIB, DivIC, FtsA, FtsL, Pbp1-3, SepF, GpsB, RodA (Steele *et al.*, 2011) and DivIVA (Bottomley *et al.*, 2017). Although some of these interactions may be false positives, it clearly suggests that EzrA is a central protein for proper cell cycle progression and cell wall synthesis in *S. aureus*. It has indeed been shown that EzrA plays a key role in staphylococcal cell size homeostasis; different levels of EzrA in the cells influence the cell size (Steele *et al.*, 2011, Jorge *et al.*, 2011). We also observed the same when *ezrA* was targeted using the CRISPRi system (Fig. 5). Furthermore, lack of EzrA causes mislocalization of other key cell division proteins such as FtsZ, GpsB and PBPs (Jorge *et al.*, 2011, Steele *et al.*, 2011). Thus, disrupting the localization or functionality of EzrA, which may be the case in cells lacking CozEa and CozEb, will therefore likely have large pleiotropic effects on different cell cycle processes, and is consistent with the results of the current study. Note, however, that there are conflicting results in the literature regarding the essentiality of *ezrA* (Steele *et al.*, 2011, Jorge *et al.*, 2011). The results from our CRISPRi depletion suggest that *ezrA* is non-essential for growth under our experimental conditions. Thus, it is likely that CozEa/CozEb have other roles in *S. aureus* yet to be identified. Since the CozEa-EzrA and CozEb-EzrA interactions were found by testing a collection of proteins in a heterologous bacterial two-hybrid assay, there may be important CozEa/CozEb interaction partners that we have not yet identified. It also remains to be determined whether the abnormal nucleoid-staining pattern is directly affected by CozEa/CozEb, or if this is an indirect effect of the compromised cell division control.

Our results also show that the influence of CozE on cell division observed in *S. aureus* was conserved in ovococcal *S. pneumoniae*. Just like in *S. aureus*, CozE^Spn^ could interact with EzrA^Spn^ in bacterial two-hybrid assays and depletion of CozE^Spn^ in *S. pneumoniae* caused aberrant cell division placement as observed by mislocalization of EzrA^Spn^-GFP. In line with this, EzrA^Spn^ interacts with FtsZ^Spn^, GpsB^Spn^ and DivIVA^Spn^ and is important for coordination of septal and peripheral cell wall synthesis in ovococcal *S. pneumoniae* cells (Rued *et al.*, 2017, Fleurie *et al.*, 2014b). Depletion of *ezrA*^*Spn*^ expression in *S. pneumoniae* (Fig. 5) also resulted in cells with variable sizes and nucleoid staining pattern as well as multiple or misplaced septa (Liu *et al.*, 2017). Notably, both *cozEa* and *cozEb* could complement the Δ*cozE*^*Spn*^ in *S. pneumoniae*, although the functionality of the staphylococcal proteins was reduced compared to the native CozE^Spn^.

CozE^Spn^ was identified as an essential regulator of cell elongation in *S. pneumoniae*, working through interactions with the MreCD^Spn^ and PBP1a^Spn^ (Fenton *et al.*, 2016). *S. aureus* also appears to elongate slightly during the cell cycle, but little is known about this and a machinery for peripheral peptidoglycan synthesis is lacking (Monteiro *et al.*, 2015). The results presented here suggest that CozE proteins in bacteria have additional functionalities to what was found for CozE^Spn^ and that they may act at an earlier stage of cell division to mediate proper spatial and temporal control. During the bacterial cell cycle, DNA replication, chromosome segregation, cell division and cell wall synthesis need to be coordinated spatially and temporally. The uncontrolled cell division occurring in *S. aureus* cells lacking CozEa and CozEb, may thus result in the pleiotropic effects we observed, such as aberrant chromosome replication/segregation, cell lysis and variable cell sizes. Future studies are required to unravel in more detail the molecular mode of action by which CozE proteins work. The CozE-mediated control of cell division seems to be a conserved feature between spherical *S. aureus* and ovococcal *S. pneumoniae*. Since genes encoding these proteins are widespread and found in bacteria with different cellular morphologies (Fenton *et al.*, 2016), it will be interesting to unravel how CozE proteins function in bacterial cells of various shapes.

## Experimental procedures

### Bacterial strains, growth conditions and transformation

Bacterial strains used in this study are listed in S2 Table. *S. aureus* was routinely grown at 37°C in brain-heart-infusion BHI broth with shaking or on BHI agar plates at 37°C. When appropriate, 5 μg/ml erythromycin, 10 μg/ml chloramphenicol or 100 μg/ml spectinomycin was added for selection. For induction of gene expression, different concentrations of IPTG was added. *S. pneumoniae* was grown in C medium (Lacks & Hotchkiss, 1960) at 37°C without shaking or on Todd-Hewitt (TH) agar plates at 37°C. When appropriate, 400 μg/ml kanamycin, 200 μg/ml streptomycin or 100 μg/ml spectinomycin was added to the growth medium for selection. *Escherichia coli* was grown at 37°C in LB medium with shaking or on LA plates at 37°C with 100 μg/ml ampicillin or 50 μg/ml kanamycin added for selection.

Transformation of *E. coli* was performed with a standard heat shock protocol. *S. aureus* was transformed with electroporation using plasmid DNA isolated from *E. coli* DC10B (Monk *et al.*, 2012) or IM08B (Monk *et al.*, 2015). Preparation of electrocompetent cells and electroporation were performed essentially as described before (Lofblom *et al.*, 2007). Constructs were introduced into *S. pneumoniae* using natural transformation as described before (Stamsås *et al.*, 2017).

### Construction of plasmids for the CRISPRi system

Construction of plasmid pLOW-dCas9. The *dcas9* gene was amplified from plasmid pJWV102-dcas9 (Liu *et al.*, 2017) using primers mk41 and mk42. The fragment and the vector pLOW-ftsZ-m(sf)gfp were both digested with SalI and NotI and ligated to produce the pLOW-dCas9 construct where *dcas9* is placed downstream of an IPTG-inducible promoter. The ligation was transformed into *E. coli* IM08B with ampicillin selection and correct construct was verified by PCR and sequencing. All plasmids in this study are listed in S3 Table, while all primers are listed in S4 Table.

Constructions of plasmids expressing single guide RNA. The single guide RNA (sgRNA) construct, containing a transcriptionally isolated sgRNA (see Fig. 2) driven by a constitutive promoter, was cut out from vector pPEPX-sgRNA(*luc*) (Liu *et al.*, 2017) using PstI and BamHI. The fragment was ligated into the corresponding sites of vector pCG248 (Helle *et al.*, 2011) (thus removing the *xyl*/*tet* regulatory system from this vector) to produce pCG248-sgRNA(*luc*). The construct was verified by sequencing.

New sgRNAs were then made by replacing the 20 nt base-pairing region with an inverse PCR approach; using the pCG248-sgRNA(*luc*) as template, new sgRNA-plasmids were amplified using one reverse phosphorylated primer mk200 annealing immediately upstream of the sgRNA, combined with a gene-specific forward primer containing the base-pairing region as overhangs. The product was treated with DpnI to remove the template plasmid and ligated using T4 DNA ligase prior to transformation into *E. coli* IM08B with ampicillin selection. Constructs were verified by sequencing. The resulting plasmids were named pCG248-sgRNA(x), where x denotes the name of the gene to be targeted. Selection of the gene-specific base pairing region to be used was done using established criteria (Liu *et al.*, 2017, Peters *et al.*, 2016).

For construction of the double-sgRNAs targeting both *cozEa* and *cozEb*, the fragment containing the sgRNA(*cozEb*) was cut out from the plasmid pCG248-sgRNA(*cozEb*) using restriction sites PstI and BamHI. The resulting fragment was ligated into the PstI and BglII sites of plasmid pCG248-sgRNA(*cozEa*). The resulting plasmid, pCG248-sgRNA(*cozEa*-*cozEb*), expresses two sgRNAs targeting both *cozEa* and *cozEb*. See also Fig. S2.

CRISPR interference. In order to obtain *S. aureus* strains for CRISPRi, the plasmid pLOW-dCas9 was first introduced with erythromycin selection. Then, in a second step, the sgRNA-containing plasmid pCG248(x), was introduced with combined chloramphenicol and erythromycin selection (in order to retain both plasmids). Cells were then grown in the presence of IPTG to induce expression of *dcas9*.

### *S. aureus* plasmid and strain construction

Construction of strain with constitutive GFP expression. A fragment containing a spectinomycin resistance gene and a gene encoding a monomeric superfolder GFP, *m(sf)gfp*, was first assembled by overlap extension PCR. The spectinomycin resistance cassette was amplified from pCN55 (Charpentier *et al.*, 2004) using primers mk203 and mk204. The *m(sf)gfp* gene was amplified from plasmid pMK17 (Kjos *et al.*, 2016) using primer im84 and im2. The primers mk204 and im84 contain overlapping sequence, and the *spc-m(sf)gfp* fragment could then be assembled in a second amplification step with outer primers mk203 and im2. The resulting fragment contains EcoRI sites on both ends introduced by overhangs in the primers. The fragment was digested with EcoRI and ligated into the corresponding site of plasmid pMAD-int2-luc. The ligation was transformed in *E. coli* IM08B with ampicillin selection. The resulting construct, pMAD-int2-luc-spc-gfp, was verified by PCR and sequencing. The temperature sensitive pMAD-derivative vector (Arnaud *et al.*, 2004) was transformed in *S. aureus* RN4220 at 30°C with erythromycin and X-gal selection. Integration of the plasmid into the chromosome and excision to construct the integration of P3-luc-spc-gfp in the *int*-locus (Fagerlund *et al.*, 2014) was performed as described (Arnaud *et al.*, 2004) with spectinomycin selection.

Construction of *ΔcozEa∷spc* and *ΔcozEb∷spc*. Vectors for deletion of *cozEa* and *cozEb* were made in pMAD. The constructions *cozEa∷spc* and *cozEb∷spc* were first assembled by overlap extension PCR as follows: The spectinomycin resistance cassette (*spc*) was amplified from plasmid pCN55 using primers mk188 and mk189. The *cozEa* upstream region was amplified with primers mk182 and mk184 and the downstream fragment with primers mk185 and mk187. The three fragments were assembled using overlap extension PCR and amplified using the outer primers mk183 and mk186. The outer primers contain restriction sites for NcoI and BamHI, and the cozEa_up – spc – cozEa_down fragment was ligated into the NcoI and BamHI sites of pMAD. The ligation was transformed into *E. coli* IM08B and correct transformants containing the pMAD-*cozEa*∷*spc* plasmid were verified by PCR and sequencing.

pMAD-*cozEb*∷*spc* plasmid was constructed in a similar way. The *spc* fragment was amplified in the same manner as above. The *cozEb* up- and downstream regions were amplified using primers mk190 and mk192, and mk193 and mk195, respectively. The resulting fragments were fused by overlap extension PCR using primers mk191 and mk194, and the resulting fragment (cozEb_up – spc – cozEb_down), was ligated into the NcoI and BamHI sites of pMAD.

Finally, the pMAD-*cozEa*∷*cam* plasmid was constructed by amplifying the upstream region with primers mk183 and mk259 and the downstream region with primers mk260 and mk186. A chloramphenicol resistance cassette was amplified from plasmid pRAB11 (Helle *et al.*, 2011), using primers mk257 and mk258. The fragments were fused by overlap extension PCR and ligated into the NcoI and BamHI sites of pMAD.

Construction of the deletion strains was done as previously described for the temperature sensitive pMAD system (Arnaud *et al.*, 2004). Briefly, the plasmids were transformed into *S. aureus* SH1000 with erythromycin selection with incubation at permissive temperature of 30°C. X-gal was also added to the transformation plates and blue colonies were re-streaked once at 30°C. One colony was then picked and grown in medium without selection at 30°C for 2 hours before the tube was transferred to non-permissive temperature for plasmid replication (43°C) for 6 hours. The culture was then plated on TSA with spectinomycin and X-gal at 43°C. White colonies, where double crossover had taken place to replace the gene of interest with the spectinomycin cassette were re-streaked on two separate plates to verify that they were spectinomycin resistant and erythromycin sensitive. Correct constructs were further verified by PCR and sequencing. The Δ*cozEa*∷*spc* deletion strain was named SAMK24 and the Δ*cozEb*∷*spc* deletion strain SAMK21.

<u>Construction of pLOW-*cozEa-m(sf)gfp* and pLOW-*cozEb*-*m(sf)gfp*</u>. *m(sf)gfp*, was first inserted into the plasmid pLOW-FtsZ-GFP (Liew *et al.*, 2011) (replacing the *gfp* gene). The *m(sf)gfp* gene with linker was amplified from plasmid pMK17 (Kjos *et al.*, 2016) using primers im1 and im2 and ligated into the BamHI and EcoRI sites of plasmid pLOW-FtsZ-GFP. The resulting construct, pLOW-*ftsZ*-*m(sf)gfp*, was verified by PCR and sequencing. To construct pLOW-*cozEa-m(sf)gfp* and pLOW-*cozEb-m(sf)gfp, ftsZ* was replaced with *cozEa* or *cozEb* in this vector. *cozEa* was amplified using primers im10 and im11, while *cozEb* was amplified using primers im12 and im13, both using genomic DNA from SH1000 as template. The fragments were digested with SalI and BamHI and ligated into the respective sites of vector pLOW-ftsZ-m(sf)gfp. The constructs were verified by PCR and sequencing.

### Strain construction for S. pneumoniae

*Construction of P*_*comX*_*-cozE*^*Spn*^, *P*_*comX*_*-cozEa, P*_*comX*_*-cozEb and deletion of cozE*^*Spn*^. The ectopic P_*comX*_*-cozE*^*Spn*^ construct integrated in the *cpsO-cpnN* locus of *S. pneumoniae* has been described previously (Straume *et al.*, 2017).

For construction of P_*comX*_-*cozEa* and P_*comX*_-*cozEb*, primers gs693/gs694 were used to amplify the *cozEa* gene and primers GS691/GS692 were used to amplify the *cozEb* gene, both using genomic DNA from *S. aureus* SH1000 as template. Using strain *S. pneumoniae* SPH131 as template, the *P*_*comX*_ and 800 bp upstream region in the *cpsO-cpsN* locus were amplified with primers khb31/khb33 and the *cpsO-cpsN* downstream fragment was amplified with primers khb34/khb36. The three fragments contain overlapping sequences introduced in the primers, and they were assembled by overlap extension PCR to create P_comX_-*cozEa* and P_comX_-*cozEb*. The constructs were transformed into strain SPH131 (containing a Janus cassette in the *cpsO*-*cpsN* locus) and transformants were selected on plates with streptomycin. The resulting strains were named GS1169 and GS1170.

The native pneumococcal *cozE*^*Spn*^ (*spr0777*) gene was replaced with a Janus cassette in strains GS1169, GS1170 and KHB432 as described before (Straume *et al.*, 2017). Since *spr0777* is essential, different concentrations of the transcription inducer ComS (0, 0.2 and 2 μM) were added during all transformation steps to induce expression of the various *cozE* genes from the P_*comX*_ promoter. Transformants were selected on plates containing kanamycin. The number of colonies were counted and the transformants were screened for the presence of the pneumococcal *cozE*^*Spn*^ gene with primers gs337 and gs338 for each ComS concentration.

Construction of *ezrA*^*Spn*^*-yfp*. An *ezrA-yfp_spc* fragment was assembled by overlap extension PCR. The ezrA_up fragment was amplified from *S. pneumoniae* R6 using primers mk288 and mk289, while the ezrA_down fragment was amplified using primers mk292 and mk293. The *yfp_spc* fragment was amplified from strain MK123 using primers mk290 and mk291. Due to overhangs in the primers, the three fragments could be assembled using outer primers mk301 and mk302, to produce the *ezrA-yfp_spc* fragment, which integrates in the pneumococcal chromosome to replace the native *ezrA* gene with an *ezrA-yfp* fusion gene. The fragment was transformed into *S. pneumoniae* and transformants were selected on plates with spectinomycin. Correct transformants were verified by PCR.

### Total RNA isolation, cDNA synthesis and qPCR

Overnight cultures were diluted to OD_600_ = 0.05 in 20 ml BHI containing 10 μg/ml chloramphenicol and 5 μg/ml erythromycin. There were two cultures of each strain, one of them was induced with 150 μM IPTG. Cells were harvested from 10 ml culture at OD_600_ = 0.4 by centrifugation at 4000 × *g* at 4°C for one minute, and the pellets were immediately frozen in liquid nitrogen. The cells were lysed by mechanical disruption in Lysing Matrix B, 2 mL tubes (MP Biomedicals) by FastPrep®-24 (MP Biomedicals). The disruption was done at maximum speed for 3 × 20 seconds, with cooling on ice between the runs. Total RNA was extracted using RNeasy Mini kit following the manufacturers’ description (Qiagen). Eluted RNA was treated with DNase I for removal of residual DNA, following the description of the manufacturer (Invitrogen). Thereafter, DNase was removed by Phenol-chloroform extraction. cDNA was synthesized using Superscript^™^ III Reverse Transcriptase (Invitrogen). Twenty-five ng cDNA was used as template for qPCR performed with PowerUp^™^ SYBR^™^ Green Master Mix (Applied Biosystems) in a StepOne Plus machine (Applied Biosystems). The setup included triplicates for each of the target genes for every sample. Primers im126 and im127 were used to target the reference gene, *pta* (Valihrach & Demnerova, 2012). Primers im130 and im131 were used to target *cozEa*, and primers im132 and im133 to target *cozEb*. The differential expression of *cozEa* and *cozEb* between non-induced and induced conditions was calculated according to the Pfaffl-method (Pfaffl, 2001).

### Bacterial two-hybrid analysis

Construction of plasmids. Genes of interest were fused in frame to either 5` end or 3` end of either the T18 or the T25 domain of adenylate cyclase from *Bordetella pertussis* using the four vectors (pKT25, pKNT25, pUT18, pUT18C) provided by the manufacturer (Euromedex). Primers used for amplification of the genes are listed in S4 Table. The amplified fragments were digested (restriction sites indicated in the S4 Table) and ligated into the corresponding restriction sites in the vectors. Ligations were transformed into *E. coli* XL1-Blue cells, and selected on 1 % glucose LA plates containing either 50 μg/ml kanamycin or 100 μg/ml ampicillin. Correct plasmids were verified by PCR and sequencing.

Bacterial two-hybrid assay. Bacterial two-hybrid assays (Karimova *et al.*, 2005) were performed as described by the manufacturer (Euromedex). Briefly, two plasmids, one containing a fusion to the T18 domain and the other a fusion to the T25 domain, were co-transformed into *E. coli* BTH101. The transformants were selected on LA plates containing 50 μg/ml kanamycin and 100 μg/ml ampicillin for selection. Five random colonies were picked per assay and grown in liquid LB containing kanamycin and ampicillin to OD_600_ 0.3, before 2.5 μl of the cell culture was spotted on LA plates supplemented with 50 μg/ml kanamycin, 100 μg/ml ampicillin, 40 μg/ml of X-gal and 0.5 mM IPTG. The plates were incubated protected from light at 30°C for 20 h to 48 h. Positive interactions are indicated by appearance of blue colonies, while white colonies indicate no interaction. All interaction assays were repeated with at least five independent replicates.

### Isolation of peptidoglycan and HPLC-analysis

Strains GS1167 and SAMK15 were inoculated in 60 ml BHI containing 10 μg/ml chloramphenicol and 5 μg/ml erythromycin. At OD_600_ 0.2 these cells were transferred to 1.5 liters of BHI containing 10 μg/ml chloramphenicol, 5 μg/ml erythromycin and 150 μM IPTG. When reaching OD_600_ = 0.3, cells were harvested at 8000 x *g* for 10 minutes. Peptidoglycan was isolated according to the protocol described by Vollmer (Vollmer, 2007). The isolated peptidoglycan was lyophilized and resuspended in water to a final concentration of 50 mg/ml.

HPLC analysis of muropeptides was performed as described by Vollmer (Vollmer, 2007) and Carvalho *et al* (Carvalho *et al.*, 2015) with minor changes. Briefly, to remove cell wall teichoic acids, ten milligrams of purified peptidoglycan were treated with 1.5 ml 48 % HF at 4°C for 48 hours with gentle mixing. The HF-treated peptidoglycan was collected by centrifugation at 20 000 x *g* for 30 minutes and washed two times with 1.5 ml of dH_2_O, once with 1.5 ml of 50 mM Tris-HCl (pH 7.4) and finally twice with 1.5 ml of dH_2_O. One mg of HF-treated peptidoglycan was digested with 5000 U mutanolysin at 37°C for 18-20 hours in a final volume of 100 μl containing 12.5 mM NaH_2_PO_4_ (pH 5.5). The sample was boiled for 20 minutes before insoluble material was removed by centrifugation at 20 000 *x g* for 30 minutes. The supernatant was added with 0.5 M Na-borate pH 9.0 (1:1 volume) and treated with 1-2 mg of Na-borohydride for 30 minutes at room temperature to reduce the sugars. The reaction was stopped by adjusting the pH to 2.0 using 20 % phosphoric acid. Muropeptides were separated on a C18 column (Vydac 218TP C18 5 mm, Grace Davison Discovery Sciences) at 52°C using a linear 155-minutes gradient of methanol from 5-30 % in 0.1 M NaH_2_PO_4_ (pH 2.0) at a flow rate of 0.5 ml/min. Eluted muropeptides were detected at 206 nm.

### Phase contrast and fluorescence microscopy

Microscopy was performed on a Zeiss AxioObserver with ZEN Blue software. Images were captured with an ORCA-Flash4.0 V2 Digital CMOS camera (Hamamatsu Photonics) through a 100x PC objective. For fluorescence microscopy, HPX 120 Illuminator (Zeiss) was used as a light source. Image analysis was performed using MicrobeJ (Ducret *et al.*, 2016) and plotting was done in RStudio.

### Transmission electron microscopy

Strains SH1000, SAMK21 and SAMK24 were grown to OD_600_ = 0.4 prior to sample preparation. GS1167 and SAMK15 were pre-grown to OD_600_ = 0.1, after which the cultures were diluted 64-fold in medium with or without 150 μg/ml IPTG and grown until OD_600_ = 0.3. Cells were fixed by adding a solution of 4 % parafomaldehyd (w/v) and 5 % glutardialdehyd (w/v) in 1 x PBS pH 7.4 to the cell culture in a 1:1 ratio. The fixation mix was incubated 1 hour in room temperature and kept overnight at 4 °C. The next day the cells were washed three times in PBS and three times in cacodylate buffer (CaCo) before being post-fixed for one hour in 1 % OsO_4_ in 0.1 M CaCo. Cells were washed three times in CaCo buffer, infiltrated in 3 % agarose and washed again three times in CaCo buffer. The samples were then dehydrated in a gradient series of 70 %, 90 %, 96 %, and 100 % ethanol (15 min for each ethanol concentration). Infiltration in LR White resin was then performed in multiple steps; LR White resin:EtOH in a ratio 1:3 was first incubated overnight, then a ratio of 1:1 for 7 hours, a ratio of 3:1 overnight and finally 100 % LR White resin overnight. Then the samples were embedded in 100 % LR White resin at 60°C for 72 hours. Thin sections were made and stained with uranyl acetate and potassium permanganate. The samples were analyzed in a FEI MORGAGNI 268 electron microscope.

### Growth assays

Growth assays were performed in a Synergy H1 Hybrid Reader (BioTek) microtiter plate reader at 37°C. Five ml of cell culture were grown to exponential phase, OD_600_ = 0.4 before being harvested, resuspended in fresh BHI medium and diluted to OD_600_ = 0.05. Appropriate antibiotics were always present. Each well in the microtiter plate was added 280 μl diluted cell culture. IPTG (150 μM) was added to the wells when appropriate. Measurements of OD_600_ were taken every 10^th^ minute throughout growth.

### Genome resequencing and analysis

Genomic DNA was isolated from *S. aureus* SH1000, SAMK21 and SAMK24 using the NucleoBond AXG 100 kit (Macherey-Nagel). For *S. aureus* SH1000, library for sequencing was created using the Nextera XT DNA library preparation kit (Illumina), and the sequencing was performed using an in-house Illumina MiSeq. For SAM21 and SAMK24, PCR-free library preparation and sequencing (HiSeq4000 PE151) was performed by BGI Hong Kong. Sequences assembly to the *S. aureus* NCTC8325 reference genome and SNP detection were done using Geneious version 10.1 (Kearse *et al.*, 2012).

## Acknowledgements

We would like to thank Lene C. Hermansen at the Imaging Center, NMBU, for help with transmission electron microscopy, Davide Porcellato, NMBU, for help with genome sequencing and Simon J. Foster and Katarzyna Wacnik, University of Sheffield, for providing the strain SH4639 (SH1000, *ezrA-gfp*). The work was funded by the Research Council of Norway (www.forskningsradet.no, grant number 250976 awarded to M.K).

## Author contributions

I.M, J.W.V, L.S.H. and M.K. conceived the study. G.A.S., I.M. D.S., Z.S. and M.K. performed experiments. G.A.S, I.M. and M.K. wrote the manuscript. D.S., J.W.V., L.S.H. edited the manuscript.

